# Predictive coordination of breathing during speaking and listening

**DOI:** 10.1101/2022.11.23.517631

**Authors:** Omid Abbasi, Daniel S. Kluger, Nikos Chalas, Nadine Steingräber, Lars Meyer, Joachim Gross

## Abstract

It has long been known that human breathing is altered during listening and speaking compared to rest. Theoretical models of human communication suggest two distinct phenomena during speaking and listening: During speaking, inhalation depth is adjusted to the air volume required for the upcoming utterance. During listening, inhalation is temporally aligned to inhalation of the speaker. While evidence for the former is relatively strong, it is virtually absent for the latter. We address both questions using recordings of speech envelope and respiration in 30 participants during 14 minutes of speaking and listening. We extend existing evidence for the first phenomenon by using the speech envelope to show that inhalation depth is positively correlated with the total power of the speech envelope in the following utterance. Pertaining to the second phenomenon, we provide first evidence that inhalation during listening to your own speech is significantly more likely at time points of inhalation during speaking. These findings are compatible with models that postulate alignment of internal forward models of interlocutors with the aim to facilitate communication.

## Introduction

Human speech production fundamentally relies on respiration which—in turn—is critical for preserving homeostasis. It is therefore noteworthy that during speaking the stereotypical rhythmic breathing process changes: Breathing during speaking is more variable in peak inhalation amplitude and breathing rate; it is also characterized by a more asymmetric pattern of short inhalations and long exhalations (Fuchs and Rochet-Capellan, 2021). However, there is a practical upper limit to the duration of a respiration cycle during speaking (Grosjean et al., 1979; Pierrehumbert, 1979). Supplying the body with sufficient oxygen as well as the capacity of the lung to supply sufficient air pressure for the articulators to resonate and produce speech sounds determine this limit individually.

Within this limit, breathing in conversation can be affected by cognitive factors of the speech production process. Here, we focus on two putative mechanisms for controlling breathing in conversations that both rely on predictions—one in speaking and another in listening. Both these mechanisms were put forward in a recent active inference model by Friston & Frith (2015) which describes human communication as two dynamic systems that are coupled via sensory information and aim to minimize prediction errors: A generative (forward) model that is likely supported by cerebello-thalamo-cortical connections computes predictions in the speaker and listener. In the speaker, the forward model represents predicted sensory consequences of their own speech and allows the speaker to adjust parameters like speech volume, speed, or articulation based on the proprioceptive and auditory feedback. Pertaining to speech breathing, there is evidence that the forward model also informs respiration based on upcoming utterances. This idea is supported by the fact that the higher variability of speech breathing compared to restful breathing is due to the fact that inhalation during speaking does not occur at regular intervals (as in restful breathing) but rather adapts to linguistic components in speech and is strongest at the beginning of a new utterance (Wang et al., 2010). This suggests a high level of fine control of speech breathing that requires speech planning to be tightly coordinated with breathing. Specifically, it has been proposed that during speaking, efficient speech breathing would adapt depth of inhalation at the beginning of a breath group (i.e., the words produced after a single inhalation) to provide sufficient air for the specific subsequent vocalization (Włodarczak and Heldner, 2017).

In the listener’s brain, the forward model generates predictions about the timing and content of upcoming speech. These predictions are constantly compared to incoming sensory information and updated accordingly (Arnal and Giraud, 2012). Two interrelated models (detailed below) suggest that interpersonal alignment between speaker and listener may facilitate predictions and, in turn, comprehension. If this interpersonal alignment extends to breathing, then we would expect that a listener could partly adapt their respiration to the respiratory dynamics of the speaker. The two models highlighted below provide further insights about the possible underlying mechanisms and their functional consequences.

First, the ‘interactive alignment account’ posits that in a conversation, speech production and comprehension is facilitated by alignment between interlocutors at various levels. While the original account has demonstrated alignment at every linguistic level (Pickering and Garrod, 2004) this concept also extends to temporal features in conversation such as speech rate, inter-speaker pause duration, and turn duration (Ostrand and Chodroff, 2021). Furthermore, during listening, brain areas associated with speech production are activated and likely improve comprehension (Möttönen et al., 2013; Pickering and Garrod, 2013; Watkins et al., 2003). Indeed, activity in listeners’ motor areas is at least partly temporally aligned with activity in the speaker’s motor system (Keitel et al., 2018; Park et al., 2018, 2015). This involvement of the cortical motor system in the alignment between speaker and listener could very well extend to some aspects of respiration (as a motor act) as well. As for other types of alignment, respiratory alignment could facilitate comprehension.

Second, ‘active sensing’ refers to the idea that sensory signals are not just received passively but rather actively sampled in a way that is modulated by the statistics of the received signals and internally generated predictions about the to-be-perceived signals and their current relevance (De Kock et al., 2021; Schroeder et al., 2010; Yang et al., 2018). Pertaining to this study it is interesting to note that this concept has been recently extended to encompass respiration: Animal studies have shown that respiration modulates spike rates in a variety of brain regions (Ito et al., 2014; Yanovsky et al., 2014) suggesting that dynamic brain states of cortical excitability fluctuate with the breathing rhythm. Recent evidence from non-invasive MEG work indicates that this coupling of respiratory and neural rhythms may apply to human brain function as well (Kluger & Gross, 2021; Kluger et al., 2021).

Such a respiratory alignment might be achieved by combining different sources of information. First, speech breathing can be perceived by the listener and the duration and intensity of speech breathing might indicate the duration of the subsequent exhalation (see above). Also, listeners can predict the length of a spoken sentence based on prosodic (and possibly breathing) cues (Grosjean, 1983; Lamekina and Meyer, 2022). Furthermore, it was recently shown that listeners utilize the perception of speech breathing to form temporal predictions about upcoming speech (MacIntyre and Scott, 2022). Second, the listener’s internal forward models afford predictions about the end of a turn of the speaker (Levinson and Torreira, 2015). Since inhalation happens frequently at the beginning of a sentence these predictions might be used for respiratory alignment.

In what follows, we present primary evidence for predictive coordination of breathing during speaking and listening.

## Results

Our analysis is based on latencies of peak inhalation which we extracted from 7-minute respiration time series acquired for N = 27 participants in four conditions: 1. Masked speech production (participants produced speech in the presence of white noise presented through eartubes such that they could not hear their own voice); 2. Normal speech production (without noise); 3. Listening to masked speech; 4. Listening to normal speech.

First, we assessed the breathing rate for each condition (Fig. 1a). We computed a linear mixed -effect model (LMEM) for our 2 × 2 design with the factors *condition* (speaking, listening) and *masking* (yes, no). Breathing rate was significantly faster during listening than during speaking (*t*(104) = 5.3, *p* << .001). Neither the main effect of masking nor the interaction of masking X condition were significant (*t*(104) = 1.8 and *t*(104) = 1.2; both *p* > .05). In addition, the variability (measured as standard deviation) of breathing rate was significantly higher during speaking compared to listening (*t*(104) = 2.3, *p* = .026). Again, neither the effects of masking nor the interaction of masking X condition was significant (*t*(104) = 0.3 and *t*(104) = 0.4; both *p* > .05). This pattern of results was to be expected - during speaking, respiration is constrained by the linguistic structure of the produced speech leading to longer and more variable intervals between peak inhalation (Fig. 1b + c).

**Fig. 1.**
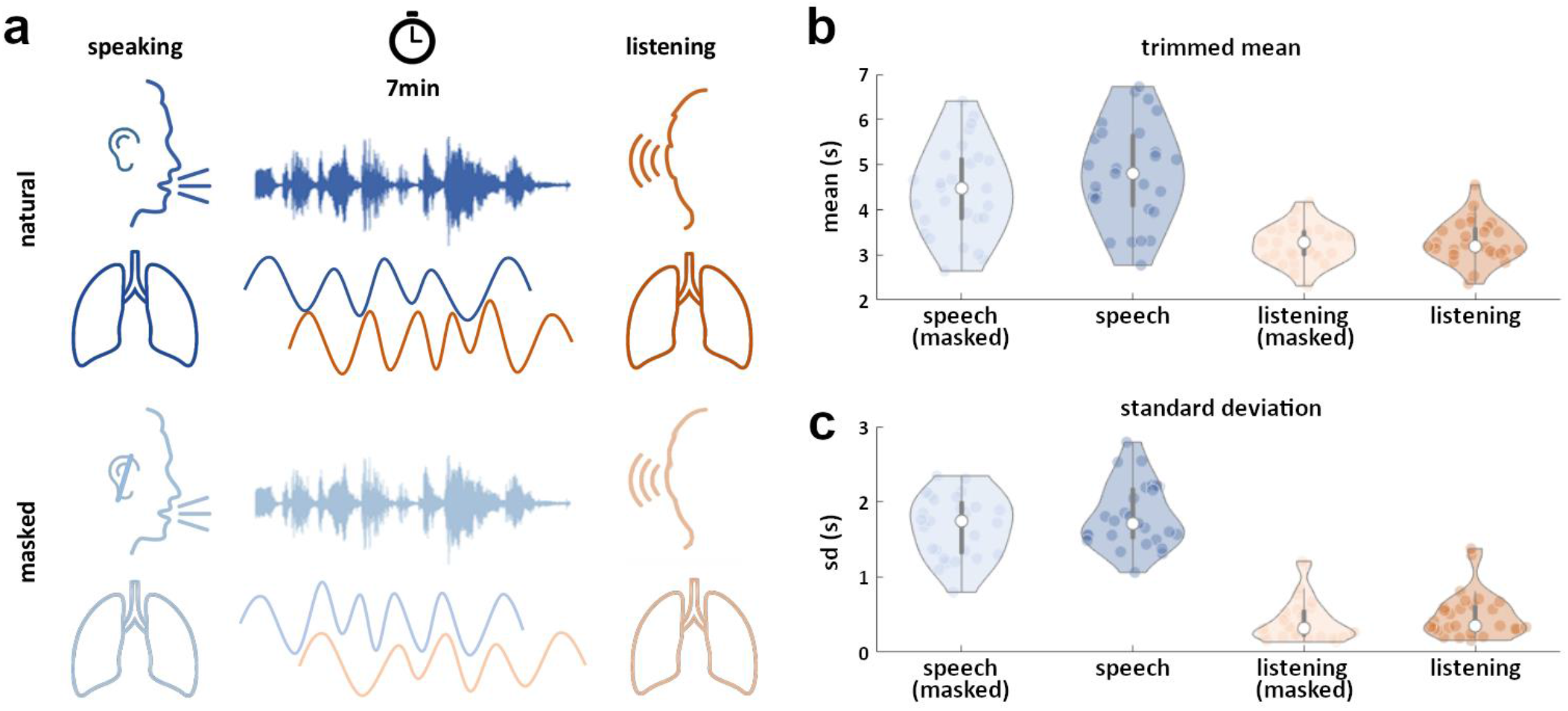
Data acquisition and respiratory cycle durations during speaking and listening. **a**, We recorded 7 minutes of respiratory data while participants were either speaking (top left) or listening to their own speech (top right). This procedure was repeated with white noise masking applied via earphones, so that participants were speaking without hearing themselves speak (*masked speech*, bottom left) and later listened to their masked speech (*masked listening*, bottom right). **b**, As expected, the LMEM model revealed significantly shorter durations of respiratory cycles for listening vs speaking (all *p* < .001, trimmed mean with 10% exclusion). **c**, Complementing the faster breathing rates during listening, the variability of cycle durations during listening was significantly reduced compared to speaking (*p* = .026).

We expected that the constraining effect of to-be-produced speech would partly determine the peak inhalation amplitude: Producing a longer or louder speech segment during one exhalation requires more air and should therefore lead to a higher peak inhalation amplitude. We tested this by employing a second LMEM to investigate the relationship between inhalation peak amplitude and the speech envelope summed over the subsequent respiration cycle (see Methods section for details). The LMEM confirmed that higher peak inhalation amplitude was significantly associated with higher total speech envelope amplitude across the subsequent respiration cycle (*t*(26) = 10.23, *p* <<0.001; see Fig. 2).

**Fig. 2:**
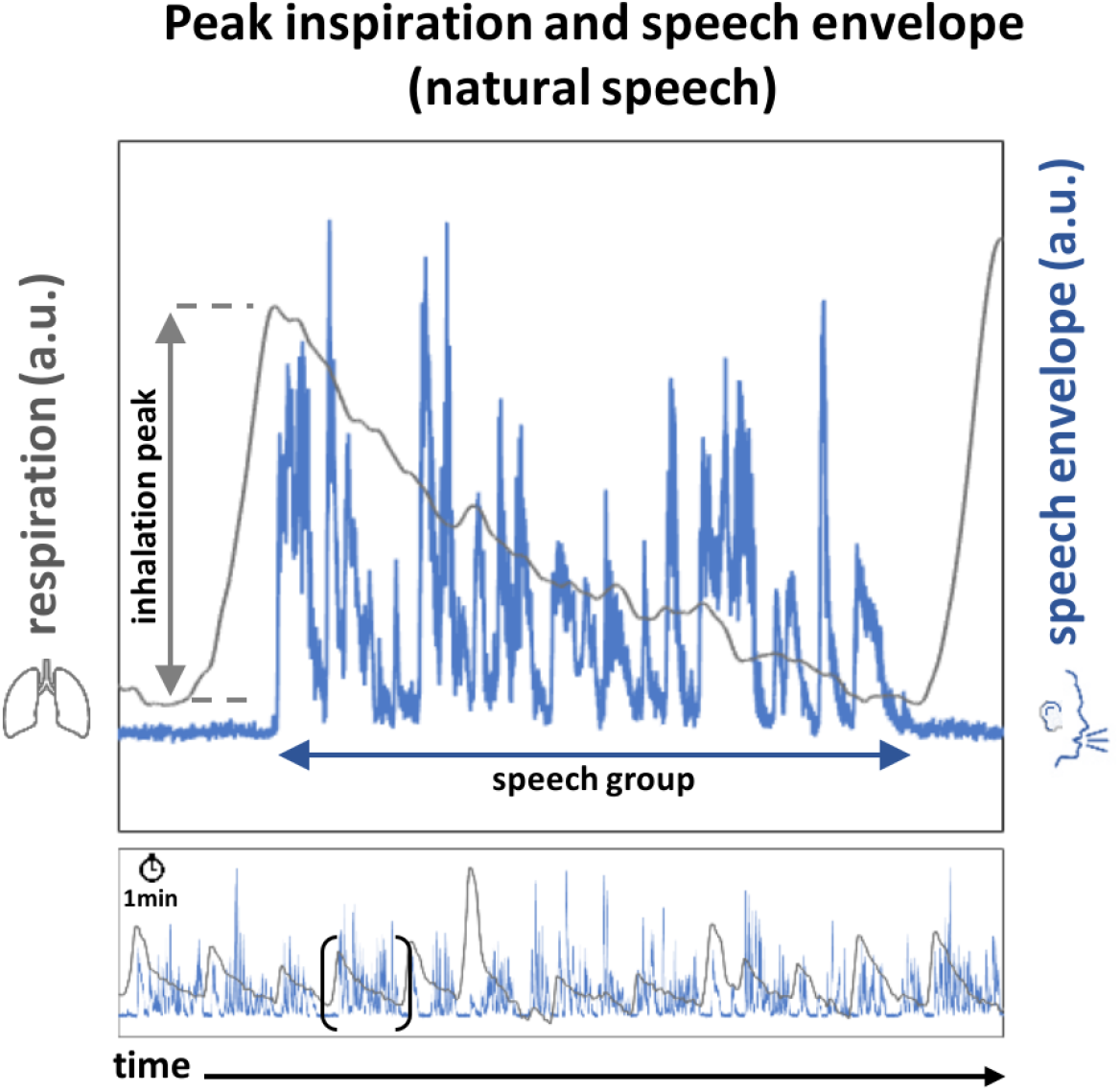
Relationship between peak inhalation and subsequent speech envelope. Exemplary respiration time course (top panel, grey line) shows the typical fast inhalation followed by slow exhalation during speaking. The blue line represents the corresponding speech envelope of this breath group. Data taken from a single 1-minute trial (see bottom panel) of a single participant.

Next, we addressed the main question of the study and tested if respiration during listening partly follows the respiration timings during speaking. Due to the significantly different breathing rates between the speaking and listening conditions we cannot expect a strong synchronization of breathing time courses where each inhalation in listening is temporally aligned to a corresponding inhalation in speaking. However, relevant events such as inhalation during listening could still have a higher probability to occur close to peak inhalation during speaking. Therefore, we identified for each inhalation peak in the speaking condition the temporally closest peak (before or after) in the corresponding listening condition and extracted the temporal peak-to-peak distance (see Fig. 3a).

**Fig. 3.**
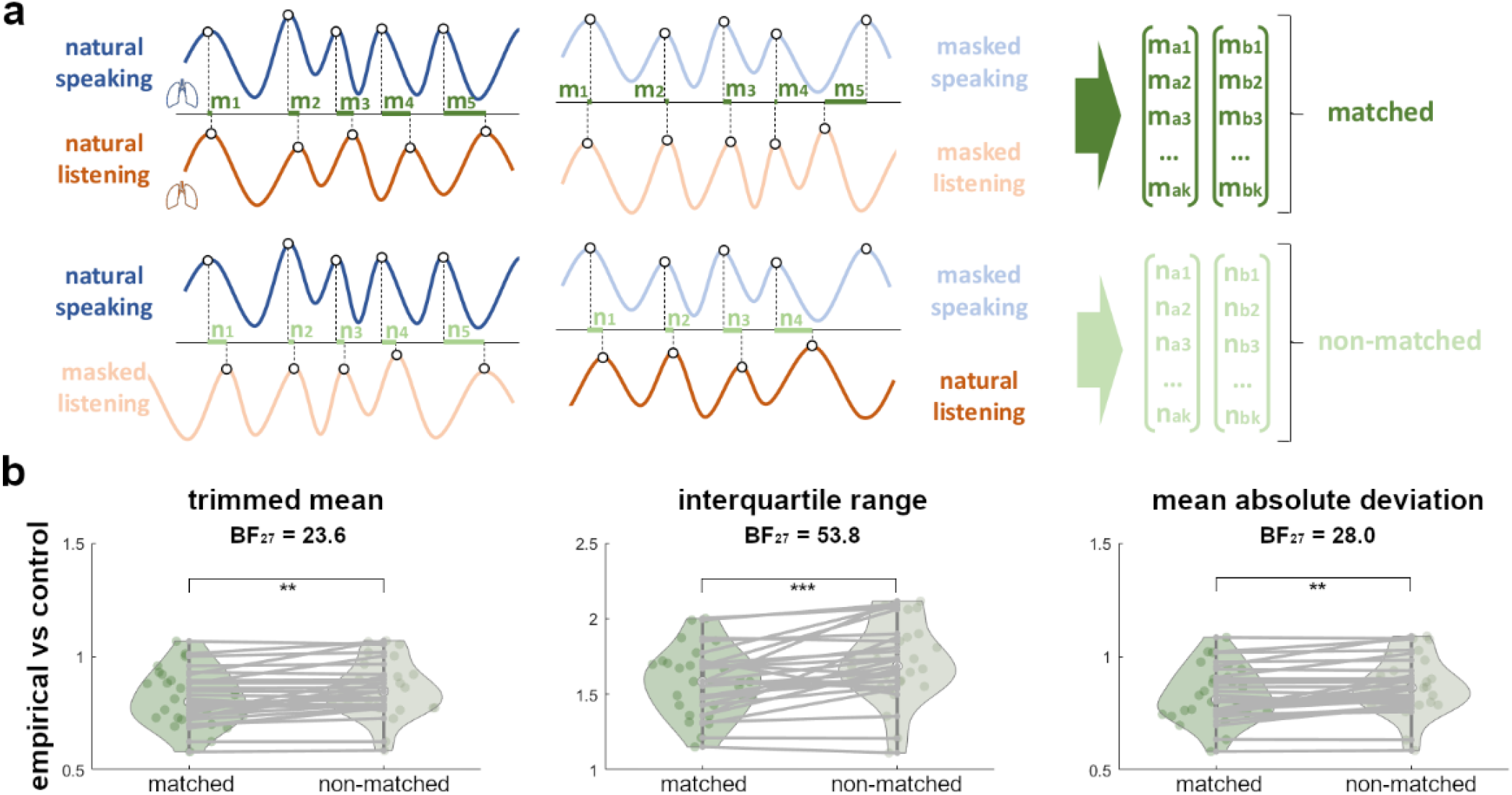
Contingencies between respiratory time courses during speaking and listening. **a**, To quantify the contingencies between breathing patterns during speaking and listening to the same speech, we computed the temporal distances between inhalation peaks in both domains: For each inhalation peak during speaking (identified by the peak detection algorithm), we computed the temporal distance to the nearest inhalation peak (before or after) in the corresponding listening condition. For the *matched* condition, we pooled first-level differences computed for [natural speaking, natural listening] and [masked speaking, masked listening] (top). For the *non-matched* condition, we computed first-level differences to the counterpart of each domain, i.e. [natural speaking, masked listening] and [masked speaking, natural listening] (bottom). **b**, Paired t-tests demonstrated a significantly closer correspondence (i.e., shorter distances) between speaking and listening for the matched (vs non-matched) condition (left panel, trimmed mean with 10% exclusion). The consistency of this decrease was corroborated by significantly lowered, variance measures like the interquartile range (middle) and mean absolute deviation (right) for matched vs non-matched speaking and listening. BF = Bayes factor. ** represents *p* < .005; *** represents *p* < .001.

Importantly, the delay between inhalation peaks during speaking and listening (which can be positive or negative) was not significantly different to 0 ms (*t*(26) = 0.07, *p* = .94), indicating that inhalation during listening is centered around inhalation latencies during speaking.

Next, we tested our main hypothesis that the temporal distance (i.e., the absolute delay) between inhalation peaks in speaking and listening is smaller than can be expected by chance. This would indicate that, during listening, participants are more likely to inhale at time points when they also inhaled during speaking.

This was tested in two ways. First, we constructed a new distribution of temporal peak-to-peak distances using *non-matched* stimuli (Fig. 3a). Specifically, we computed the temporal distance between inhalation peaks during speaking and listening of the opposite conditions (i.e., natural speaking - masked listening and masked speaking - natural listening). These distances were then compared to those within the *matched* stimuli (i.e., natural speaking - natural listening and masked speaking - masked listening). Second, we constructed an artificial sequence of inhalation time points with the same distribution of respiration cycle durations as the individual listening condition (see Methods section and Fig. 4 for design of these surrogate data). It is important to note that the mean breathing rates had a strong effect on the distribution of temporal distances of inhalation peaks. Importantly, both control distributions were specifically designed to preserve the mean breathing rate (see Methods section for more details).

**Fig. 4.**
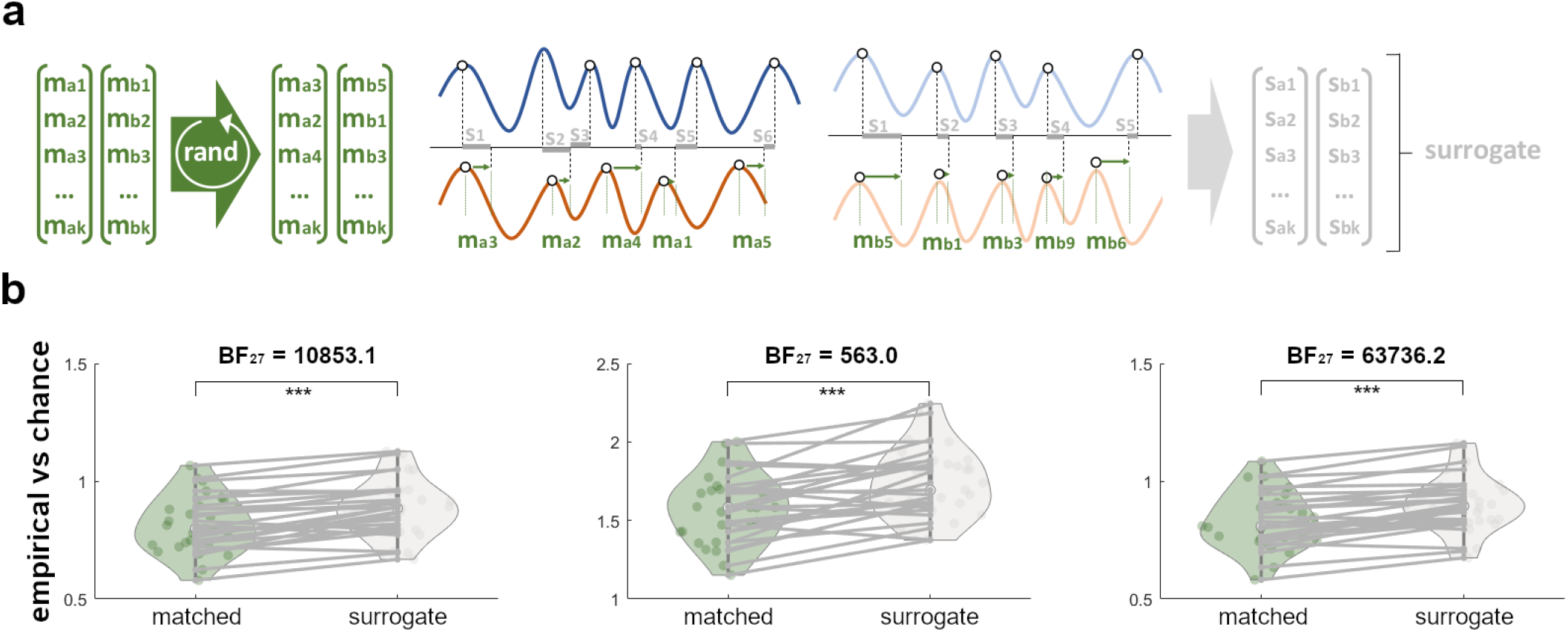
Contingency statistics against surrogate data. **a**, Individual surrogate distributions of temporal distances were constructed from the empirical distributions described in Fig. 2: For each participant, the vectors of peak-to-peak distances computed between [natural speaking, natural listening] and [masked speaking, masked listening] were shuffled separately. The elements of the resulting shuffled vectors were consecutively used to shift the empirical inhalation peaks during listening by a random (yet physiologically plausible) distance. In line with the procedure described above, we then identified the closest inhalation peak during speaking for each of the shifted time points, thus constructing vectors of surrogate peak-to-peak distances. **b**, Compared to these surrogate distributions, the empirical peak-to-peak distances were found to be significantly shorter on average (left). The consistency of this effect was indicated by a significant lowering of interquartile range (middle) and mean absolute deviation (right) for the empirical (vs surrogate) distances. BF = Bayes factor. ** represents *p* < .005; *** represents *p* < .001.

The results indicate that inhalation peaks during listening to one’s own speech were significantly closer to inhalation peaks during speaking than can be expected by chance (control 2: surrogate data; Fig. 4) or when inhalation peaks were taken from a different (non-matched) speech condition (control 1; Fig. 3).

This is true for robust estimates of the mean absolute temporal distance but also for the interquartile range as a robust measure of the spread. The effect of ‘inhalation alignment’ is subtle but highly significant, indicating that there was a significant bias (i.e., higher probability) for inhalation during listening to occur at the time of inhalation during speaking.

Finally, we investigated if, beyond the temporal alignment of respiration, the depth of inhalation was also related in speaking and listening. We tested this with an LMEM relating peak inhalation amplitude in listening to peak inhalation amplitude in speaking. The model yielded a significant relationship between depth of inhalation in speaking and listening (*t*(4373) = 2.69, *p* = .007). Taken together, our results indicate that listeners mimic not only the timing but also the depth of inhalation of their previously produced speech.

## Discussion

Our results indicate predictive coordination of respiration during speaking and listening. During speaking the peak inhalation amplitude was related to the total speech envelope summed across the breath group. The positive coefficient of the LME-model indicates that larger peak inhalation amplitude is associated with higher summed speech envelope. This means that in preparation of vocalization of a breath group speakers adapt inhalation to the air volume required for the respective breath group. Previous studies have demonstrated that peak inhalation amplitude is correlated with the duration of the subsequent utterance during reading (Fuchs et al., 2013), single sentence utterance (McFarland and Smith, 1992) and spontaneous speech (Rochet-Capellan and Fuchs, 2013a; Winkworth et al., 1995). However, the evidence is not unambiguous since in a recent study inhalation amplitude did not differ significantly between very short utterances and longer speech (Włodarczak and Heldner, 2017). We improve on previous studies by using the speech envelope instead of breath group duration only. By relating inhalation amplitude to the summed speech envelope over the subsequent breath group (instead of its duration) we are getting closer to the hypothesized mechanism underlying predictive coordination of respiration during speaking because the summed speech envelope is a better measure of speech output compared to the duration of the breath group. The required air volume for a breath group correlates with its duration but also depends on speech loudness (Huber, 2008) and more generally the speech-specific sound pressure that is adequately quantified with the speech envelope. Therefore, our results based on the speech envelope and peak inhalation amplitude further support the notion that inhalation is finely controlled based on the upcoming breath group.

While this mechanism is strikingly intuitive for energy-efficient speech production it requires very sophisticated computations. Specifically, while a speaker initiates inhalation, planning of the content of the upcoming breath group needs to be largely completed. In addition, the speaker needs a model that provides a mapping of this content to the estimated required air volume which in turn depends on loudness, the current physiological state (which is different e.g. for resting, walking, running) and other factors. And, as alluded to above, this joint speech and breath planning needs to be conducted within the constraints of the individual lungs vital capacity (i.e. volume of air available for vocalization).

The second aspect of predictive coordination of respiration studied here pertains to listening. Our results indicate that the initiation of inhalation in the listener is more likely at time points that correspond to inhalations in the speaker. This is very different to a pure 1:1 phase synchronization of respiration between speaker and listener. Such strict phase synchronization is not possible in the case of speaker-listener respiratory alignment given the very different breathing rates between speaking and listening (see Fig. 1). As a consequence there is no one-to-one mapping of inhalations between speaking and listening. Instead, our results are consistent with the idea that listeners have a preference to inhale at time points close to the inhalation of speakers. However, we like to note several caveats. First, while this partial temporal alignment is highly significant (i.e. very consistent across the group of participants) the actual effect (difference of temporal distances between real data and surrogate data; see Fig. 3b + 4b) in each individual is rather subtle. Second, in our study participants were listening to their own speech and they might have anticipated some parts during the listening condition. However, during designing our experiment, we tried to lower the chance of this anticipation by several means: Participants were measured in separate sessions for speech production and perception tasks. There were always several days’ intervals between performing these two conditions. Further, our questions were mainly about a common/general topic. Consequently, participants may not remember their answers completely. Still, it remains to be seen if our results generalize to unknown speech from a different speaker.

As outlined in the introduction, respiratory coordination between listener and speaker would be consistent with several models. All these models are to some degree based on the notion of a coupling of internal forward models of speaker and listener via sensory signals produced by the speaker’s motor system (such as sound of speech or respiration or visual cues of respiration). Therefore, in the listener parts of the motor system are aligned to the speaker possibly leading to enhanced comprehension through coordination of internal excitability states and simulation of the speaker’s internal model (see also Barsalou, 2008).

There is convincing evidence that simultaneously perceived sensory signals lead to interpersonal synchrony. Recently, Madsen and Parra performed a comprehensive study showing that watching the same movie induces intersubject correlation in participants of EEG signals, gaze position, pupil size, and heart rate, but not respiration and head movements (Rochet-Capellan and Fuchs, 2013b). In other studies interpersonal physiological synchrony has been observed for electrodermal activity and heart rate (Stuldreher et al., 2020), eye movement (Madsen et al., 2021), and for movement and respiration (Codrons et al., 2014; Paccalin and Jeannerod, 2000). Pertaining to respiration there is also plenty of evidence that it is adjusted to motor activity within an individual (Bartlett and Leiter, 2012; Rassler and Raabe, 2003; Ebert et al., 2002). Evidence for auditory–motor alignment within individuals in the context of continuous speech is however sparse and has received relatively little attention. Garssen (1979) reported that the number of respiration cycles where inhalation is aligned between speaker and listener is higher than expected by chance but only when watching a video of an actor where respiration is clearly visible and audible. However, this is a rather lax criterion of respiratory coordination and the statistics used by Garssen are incorrect. The data was compared against the mean of three instantiations of surrogate data instead of a comparison to the 95th percentile of a large number of surrogates. More recently this question was revisited by Rochet-Capellan and Fuchs (Rochet-Capellan and Fuchs, 2013b). They asked participants to listen to read speech and studied to what extent a listener aligns inhalation onset to those of the reader. Alignment was observed when listening to the female reader but not the male reader and authors concluded that findings ‘did not support stable or continuous temporal alignment of listener breathing to reader breathing.’

The absence of a continuous alignment is consistent with our results and—given the different breathing rates between speaking and listening—can be expected. However, using a different methodology and two control conditions, we find significant respiratory alignment in the sense described at the beginning of this section. Our results therefore indicate that inhalation in listeners is modulated by attended speech not only in general aspects such as breathing rate and amplitude but also in the timing, leading to a preferred inhalation of the listener near time points of inhalation in speakers. Clearly, not every inhalation in the speaker is matched with an inhalation in the listener. It therefore remains an intriguing question for further studies if the probability of speaker-listener alignment for each inhalation can be predicted—e.g. from factors such as momentary attention, emotional engagement, predictability of speakers’ breathing pattern or acoustic or linguistic aspects of listened speech.

## Acknowledgements

We acknowledge support by the Interdisciplinary Center for Clinical Research (IZKF) of the medical faculty of Münster (grant number Gro3/001/19). OA (EFRE-0400394) was supported by the EFRE. DSK (KL 3580/1-1) and JG (GR 2024/5-1; GR 2024/11-1; GR 2024/12 -1) were further supported by the DFG.

## Methods

### Participants

We recruited thirty native German-speaking participants (15 males, mean age 25.1 ± 2.8 years [M ± SD], range 20–32 years). The study was approved by the local ethics committee and conducted in accordance with the Declaration of Helsinki. Prior written informed consent was obtained before the measurement and participants received monetary compensation after their participation.

### Recording

MEG, electromyogram (EMG), respiration, and speech signals were recorded simultaneously. Only respiration and speech signals were used for this study. Details of the MEG recordings are reported elsewhere (Abbasi et al., 2022). The speech recording had a sampling rate of 44.1 kHz. Audio data was captured with a microphone which was placed at a distance of 155 cm from the participant’s mouth in order not to cause any artefacts by the microphone itself. The respiratory signal was measured as thoracic circumference by means of a respiration belt transducer (BIOPAC Systems, Goleta, USA) placed around the participant’s chest. Individual respiration time courses were visually inspected for irregular breathing patterns such as breath holds or unusual breathing frequencies, but no such artefacts were detected.

### Paradigm

Participants were asked to sit relaxed while performing the given tasks and to keep their eyes focused on a white fixation cross. This study consisted of three separate recordings: i) speech production, ii) speech production while perception of their own speech was masked, and iii) speech perception. For the speech production recording, there were seven 60-second trials for overt speech. During each trial, participants answered a given question such as ‘What does a typical weekend look like for you?’. A colour change of the fixation cross from white to blue indicated the beginning of the time period in which participants should speak and the end was marked by a colour change back to white. In the second recording, participants were asked to perform the same task as in the first recording while they heard white noise, leaving them largely unable to hear their own voice. The questions were different to the prior recording in order to prevent repetition of prefabricated answers. Questions covering neutral topics were chosen to avoid emotional confounds.

In the third recording session participants listened to audio-recordings of their own voice which were collected in the first and second recordings.

### Preprocessing and data analysis

In the preprocessing and data analysis steps, custom-made scripts in Matlab R2020 (The Mathworks, Natick, MA, USA) in combination with the Matlab-based FieldTrip toolbox (Oostenveld et al., 2011) were used in accord with current guidelines (Gross et al., 2013). Three participants where respiration recording failed were excluded from analysis.

The wideband amplitude envelope of the speech signal was computed using the method presented in (Chandrasekaran et al., 2009). Nine logarithmically spaced frequency bands between 100-10000 Hz were constructed by bandpass filtering (third-order Butterworth filters). Then, we computed the amplitude envelope for each frequency band as the absolute value of the Hilbert transform and downsampled them to 1200 Hz. Finally, we averaged them across bands and used this computed wideband amplitude envelope for all further analysis.

In all respiration signals obtained during speaking and listening we identified the time points corresponding to peak inhalation. To this end, we employed the findpeaks.m function in matlab on the z-scored time series after smoothing (Savitzky-Golay filter of order 3 and frame length 1591). Three participants were excluded because peak inhalation could not be reliably identified in their data. Results were validated by visual inspection. The temporal distance between peak inhalation times yielded the respiration cycle durations (and their variability) presented in Fig. 1. These data were subjected to a linear mixed effects model (LMEM) using the equation in Wilkinson notation:

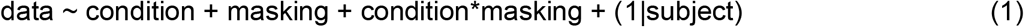

The model fit was obtained with the fitlme function in Matlab R2022a (Mathworks).

Time points of peak inhalation during speaking were used to study the relationship between peak inhalation amplitude and the summed speech envelope of the subsequent breath group (i.e. the speech envelope in the time window until next inhalation). The LMEM used the equation in Wilkinson notation:

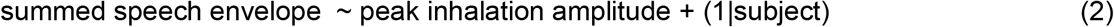

The main analysis was based on the temporal distance of inhalation peaks between listening and speaking. Recall that participants were listening to the same speech that they had themselves produced in an earlier recording session. We aimed to test if inhalation during listening was more likely to occur at time points where inhalation occurred in the speaking condition. Therefore, we identified for each inhalation peak during speaking the temporally closest inhalation peak in the listening condition. To improve statistical sensitivity we pooled the normal speech and masked speech condition leading to 14 min of data (7 min speaking, 7 min masked speaking). Finally, we statistically compared the distribution of temporal distances to two other control distributions. The first control distribution was constructed by identifying temporal distances between non-matching stimuli. While the original distribution was constructed from matched stimuli ([natural speaking, natural listening] and [masked speaking, masked listening]), the first control distribution was constructed from the non-matched stimuli pairings ([natural speaking, masked listening] and [masked speaking, natural listening]). This control distribution therefore represents the distribution of delays that can be expected by chance.

The second control distribution was constructed artificially: For each individual participant, we computed a new vector of peak inhalation times by picking a random start time for the first inhalation and then successively adding to this time point randomly picked respiration cycle durations from the individual real respiration cycle durations (see Fig. 4a). Therefore, this surrogate list had the same distribution of respiration cycle durations as the original individual listening condition - but not in the right order. This procedure would destroy any temporal alignment of inhalation between speaking and listening while preserving the overall statistics of the respiration cycle duration.

A test on the relationship between peak inhalation amplitude in speaking and listening was conducted with an LMEM using the equation:

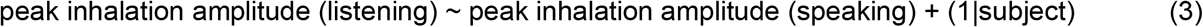

